# Common Analysis of Direct RNA SequencinG CUrrently Leads to Misidentification of 5-Methylcytosine Modifications at GCU Motifs

**DOI:** 10.1101/2023.05.03.539298

**Authors:** Kaylee J. Watson, Robin E. Bromley, Benjamin C. Sparklin, Mark T. Gasser, Tamanash Bhattacharya, Jarrett F. Lebov, Tyonna Tyson, Laura E. Teigen, Karen T. Graf, Michelle Michalski, Vincent M. Bruno, Amelia R. I. Lindsey, Richard W. Hardy, Irene L. G. Newton, Julie C. Dunning Hotopp

## Abstract

RNA modifications, such as methylation, can be detected with Oxford Nanopore Technologies direct RNA sequencing. One commonly used tool for detecting 5-methylcytosine (m^5^C) modifications is Tombo, which uses an “Alternative Model” to detect putative modifications from a single sample. We examined direct RNA sequencing data from diverse taxa including virus, bacteria, fungi, and animals. The algorithm consistently identified a 5-methylcytosine at the central position of a GCU motif. However, it also identified a 5-methylcytosine in the same motif in fully unmodified *in vitro* transcribed RNA, suggesting that this a frequent false prediction. In the absence of further validation, several published predictions of 5-methylcytosine in human coronavirus and human cerebral organoid RNA in a GCU context should be reconsidered.

**IMPORTANCE:** The detection of chemical modifications to RNA is a rapidly expanding field within epigenetics. Nanopore sequencing technology provides an attractive means of detecting these modifications directly on the RNA, but accurate modification predictions are dependent upon the software developed to interpret the sequencing results. One of these tools, Tombo, allows users to detect modifications using sequencing results from a single RNA sample. However, we find that this method falsely predicts modifications in a specific sequence context across a variety of RNA samples, including RNA that lacks modifications. Results from previous publications include predictions in human coronaviruses with this sequence context and should be reconsidered. Our results highlight the importance of using RNA modification detection tools with caution in the absence of a control RNA sample for comparison.

## INTRODUCTION

Oxford Nanopore Technologies (ONT) direct RNA sequencing (**Figure 1A**) enables detection of RNA modifications. A modified base produces an altered electrical current and/or dwell time relative to a canonical base that can be detected with algorithms (1-3).

**Figure 1.**
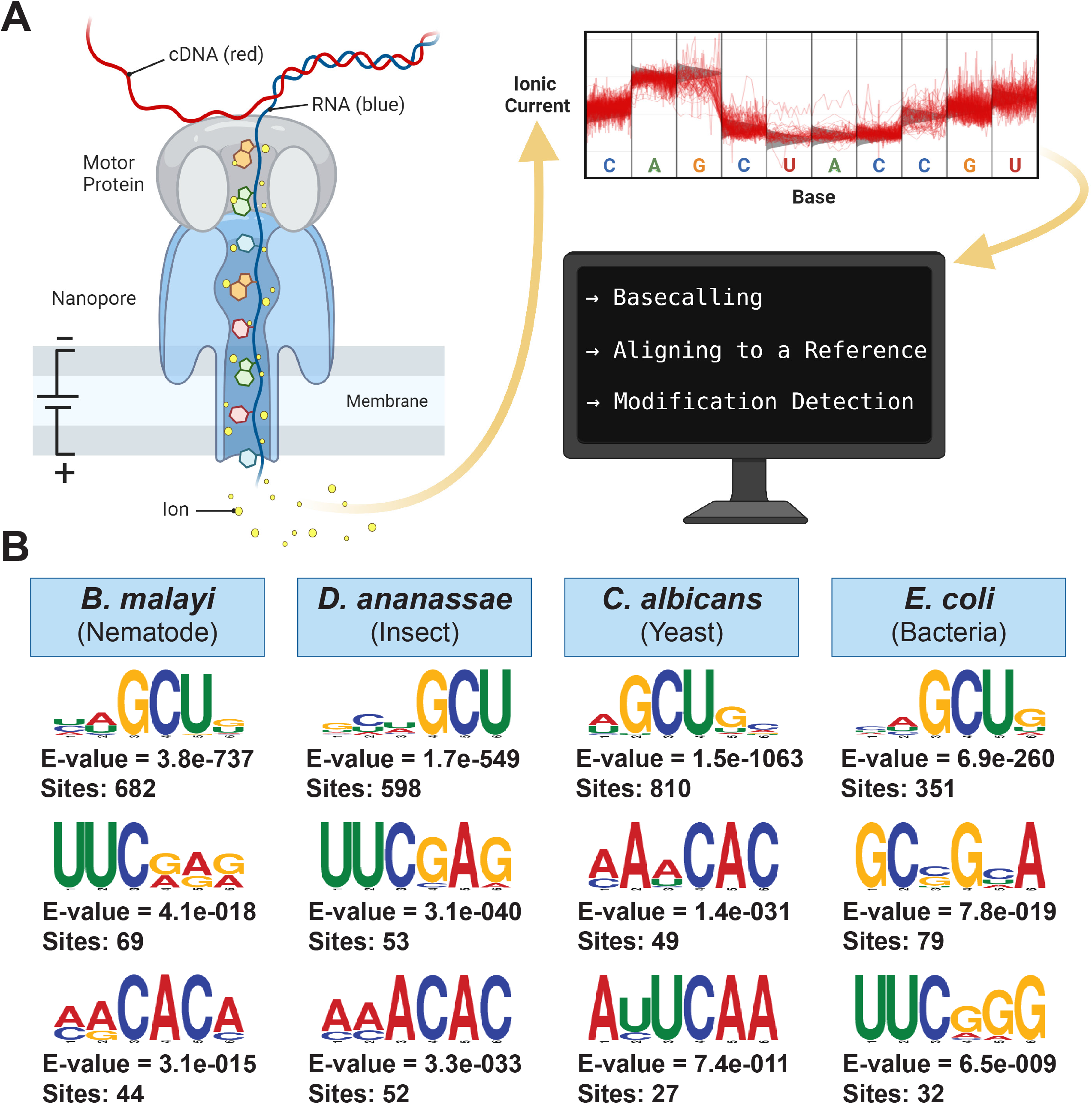
ONT direct RNA sequencing and GCU motif detection. **A** Schematic showing how RNA molecules are directly sequenced with ONT, followed by basecalling of the signal data produced by changes in ionic current, mapping to a reference, and detection of modified bases. Adapted from “Nanopore Sequencing”, by BioRender.com (2022). Retrieved from https://app.biorender.com/biorender-templates. **B** MEME Suite motifs overrepresented in the 10-nucleotide sequences surrounding the top 1000 putative modifications detected by the Tombo m^5^C “Alternative Model” for *B. malayi, D. ananassae, C. albicans*, and *E. coli*.

Tombo is a commonly used software with three methods for detecting 5-methylcytosine (m^5^C) modifications, including two for detecting m^5^C in a single sample: “Alternative Model” and “*de novo* Model”. The Alternative Model detects signals that fall within the expected range for a m^5^C modification. In contrast, the *de novo* detection method identifies differences from a canonical base model and is unable to predict the type of putative modification. Given that these methods can be run on a single sample, they have been used to predict modifications in many biological entities including *Arabidopsis thaliana* (4), SARS-CoV-2 (5), human coronavirus 229E (6), and human cerebral organoid RNA (7). Using the Alternative Model, coronavirus and the human cerebral organoid RNA are both predicted to have a modified GCU motif (6, 7).

The third Tombo method, “Sample Compare” detects modifications by comparing the raw signals of two samples but is unable to predict the type of modification. “Sample Compare” can detect differing levels of modification between two biological samples, or it can detect putative modifications in a single sample through comparisons to unmodified RNA generated by *in vitro* transcription (IVT) or knockout/knockdowns of modifying enzymes. However, this is limited by the ability to generate IVT RNAs or knockout/knockdowns. Sample Compare has been used to detect native SARS-CoV-2 modifications relative to an IVT control, but the Sample Compare predictions differed from those generated with the Alternative Model (5). Here, we looked for m^5^C RNA modifications using the Alternative Model and identified a GCU motif that is a consistent false prediction across viral, bacterial, fungal, and animal RNA.

## RESULTS AND DISCUSSION

To broadly assess RNA modification patterns, RNA was isolated from diverse organisms including *Brugia malayi* FR3 (Animal:Nematode; 90 Mbp; 14,388 genes), *Drosophila ananassae* Hawaii (Animal:Insect; 220 Mbp; 23,553 genes), *Candida albicans* SC5314 (Fungi: Saccharomycetes; 15 Mbp; 6,271 genes), and *Escherichia coli* K-12 MG1655 (Bacteria:Gammaproteobacteria; 5 Mbp; 4,723 genes) (8-11) **(Supplementary Table 1)**. The *B. malayi, D. ananassae*, and *C. albicans* RNA was prepared for ONT direct RNA sequencing with the standard protocol. Since ONT adapters require a poly(A) tail for annealing, *E. coli* total RNA was polyadenylated using *E. coli* poly(A) polymerase prior to library preparation. This resulted in a larger proportion of ribosomal RNA present in the *E. coli* data **(Supplementary Table 1)**. Sequencing resulted in 822 Mbp *B. malayi*, 809 Mbp *D. ananassae*, 2.4 Gbp *C. albicans*, and 216 Kbp *E. coli* mapped reads **(Supplementary Table 1)**.

The Tombo Alternative Model detection method was used (4-7) to detect the cytosines that deviated from the canonical model and fell within the expected range for the m^5^C model. The methylated fraction of mapped reads that were predicted to have a m^5^C was identified at each cytosine and ranked from highest to lowest. The methylated fractions range from 0 to 1.0 across the entire transcriptome, so the number of m^5^C sites cannot be predicted as these results are not binary. The 1000 most highly modified positions ranged between 0.85 and 1.0, meaning at least 85% of the reads at all 1000 positions were predicted to be modified (**Supplementary Table 2**).

Motifs associated with those cytosines that had the highest methylated fraction were detected using the MEME suite (12) on the cytosine and the 10 adjacent nucleotides. The most significant motif for all four organisms contained a GCU (**Figure 1B**) suggesting either an artifact or a m^5^C motif that spans multiple kingdoms of life, similar to the DRACH motif observed for *N*-6-methyladenosine (13-16). While the number of putative modified GCU sites varied by organism, all organisms had over four times as many putative modified GCU sites as any other motif in the top 1000 predicted modified positions.

To determine whether false predictions were occurring with the Tombo Alternative Model detection method, the 11.7 kbp Sindbis virus (SINV) RNA genome was *in vitro* transcribed and compared to native viral RNA from infected JW18 insect cells, which contains modified bases (17-20). Sequencing resulted in 320 Kbp native SINV and 1.38 Mbp of IVT SINV sequence reads (**Supplementary Table 1**), which were analyzed separately for m^5^C modifications using the Tombo “Alternative Model”. Because SINV only contains 146 GCU sites and 2,970 total cytosines, a top 1000 putative modification sites analysis is uninformative. Whereas in whole organisms, the lowest methylated fraction in the top 1000 putative modified sites is 0.85; in the virus, the lowest methylated fraction is 0.1. Conversely, if we only use sites that are at least 85% percent methylated, we would only have 14 positions for a motif analysis, which is insufficient. Therefore, instead of the motif-based analysis of the top 1000 putative modification sites, all 3-mers with a central cytosine in the viral genome were analyzed individually for the IVT and native SINV RNA. The methylated fraction in GCU motifs tended to be higher than any other 3-mer **(Figure 2A, Supplementary Figures 1-2)**.

**Figure 2.**
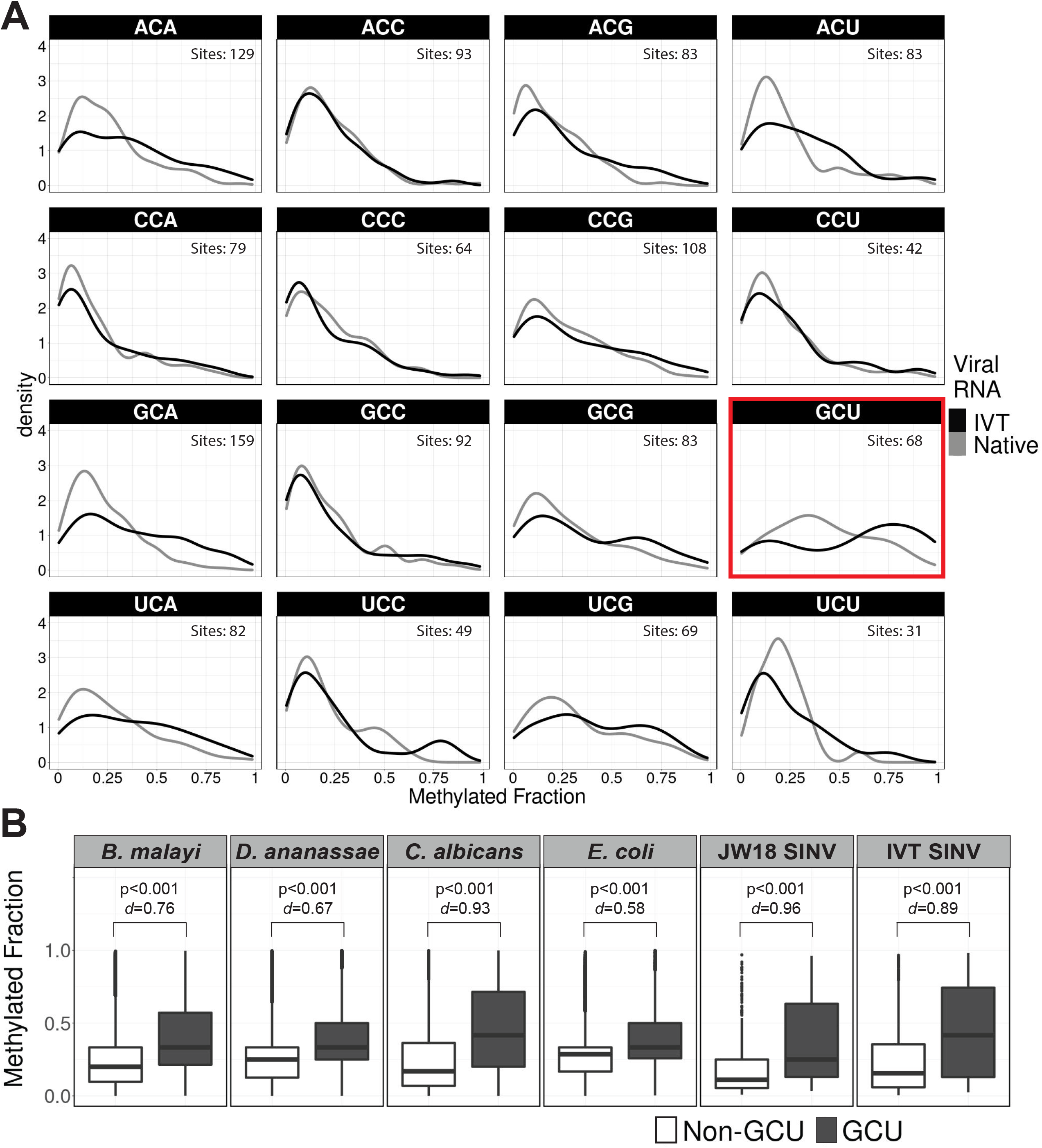
Methylated fractions predicted by Tombo “Alternative Model” for m^5^C. **A** Density plots of the methylated fraction at all 3-mers containing a central cytosine in native and IVT viral RNA. Cytosine positions were filtered for depth > 10 and methylated fraction > 0 in both IVT and native samples. Histograms available in supplemental information. **B** Boxplot showing distributions of the methylated fractions detected by Tombo m^5^C “Alternative Model.” The methylated fraction was extracted for cytosines in animal, fungal, and bacterial RNA with depth > 100 and cytosines in native and IVT viral RNA with sequencing depth > 10. Methylated fractions were grouped based on non-GCU and GCU sequence context. Statistical significance based on p-value from two-tailed Z-test and Cohen’s *d* effect size.

Furthermore, the median predicted methylated fraction for GCU sites was 1.2- to 2.7-fold higher relative to non-GCU context cytosines in *B. malayi, D. ananassae, C. albicans, E. coli*, SINV, and IVT-prepared SINV that is expected to be fully unmodified **(Figure 2B)**. The detection of “modifications” in IVT RNA indicates this is merely an artifact of the detection algorithm.

## CONCLUSIONS

The Tombo “Alternative Model” consistently overpredicts GCU as being modified in RNA from diverse taxa. This motif has been reported in numerous papers as a highly modified sequence, highlighting a need for improved methods and standards in RNA modification detection from direct RNA sequencing data. Without further experimental validation of GCU modification, results reporting GCU modifications should be considered carefully, including previously published results (6, 7).

## METHODS

Detailed methods are available in **Supplemental Information**.

### RNA isolation

*B. malayi* FR3 RNA was isolated using TRIzol (Zymo Research, Irvine, CA, USA) with tissue homogenization followed by purification with a PureLink RNA Mini column (Ambion, Austin, TX, USA) (21). Publicly available (SRR15923920) *D. ananassae* ONT direct RNA sequencing data was used (**Supplementary Table 1**) that was sequenced from RNA isolated from whole flies homogenized in liquid nitrogen using QIAzol and chloroform (22). *C. albicans* RNA was isolated using TRIzol (Zymo Research, Irvine, CA, USA) with bead beating and a PureLink RNA mini column (Ambion, Austin, TX, USA). *E. coli* RNA was isolated using an RNEasy column (Qiagen, Germantown, MD, USA) followed by polyadenylation with an *E. coli* poly(A) polymerase (NEB, Ipswich, MA, USA). Total RNA was extracted from SINV-infected JW18 cells using TRIzol reagent (Invitrogen, Waltham, MA, USA) followed by RQ1 RNase-free DNase (NEB, Ipswich, MA, USA) treatment using manufacturer’s protocol. SINV IVT RNA was generated by SP6-driven transcription with MEGAscript (Thermo Fisher Scientific Inc., Waltham, MA, USA) and the TE12 BC 4.10 plasmid, followed by lithium chloride precipitation.

### Sequence analysis

Libraries were prepared with the SQK-RNA002 Direct RNA Sequencing Kit (Oxford Nanopore Technologies, Oxford, UK) and sequenced with an R9 flow cell using an ONT MinION. Reads were basecalled using Guppy v.6.4.2 with the “rna_r9.4.1_70bps_hac.cfg” model and filtered using a minimum quality score cutoff of 7. Reads were aligned to transcriptome references using minimap2 v.2.24 (23). The number of reads sequenced and mapped were determined for each sample using SAMtools (24). SeqKit (25) was used to calculate the N50, bases sequenced, and bases mapped.

### RNA Modification Detection and Motif Discovery

5-methylcytosine RNA modifications were detected using the Tombo Alternative Model. Multi-read fast5 files were converted to single-read fast5 files using Oxford Nanopore Technologies fast5 API software and reannotated with Tombo preprocess annotate_raw_with_fastqs. The raw signal was then assigned to each base using Tombo resquiggle and modified bases were detected with Tombo detect_modifications alternative_model. The top 1000 regions were analyzed with MEME suite (12).

Results were plotted with R v3.6.3.

## DATA AVAILABILITY

The datasets generated during and/or analyzed during the current study are available in the NCBI Sequence Read Archive repository, <u>PRJNA944578</u> (**Supplementary Table 1**). All commands and scripts are available on Github: https://github.com/Dunning-Hotopp-Lab/Common-Analysis-of-Direct-RNA-Sequencing-Misidentification-of-5mC-Modifications-at-GCU-Motifs

## ACKNOWLEDGMENTS

This project was funded by federal funds from the National Science Foundation grant number EF 2025384 and the National Institute of Allergy and Infectious Diseases, National Institutes of Health, Department of Health and Human Services under grant numbers U19AI110820 and T32AI162579. The funding body had no role in the design of the study and collection, analysis, and interpretation of data and in writing the manuscript.

All animal care and use protocols were carried out in accordance with the relevant guidelines and regulations and were approved by the UWO IACUC, protocol 246-R4.

Author contributions are described using the controlled vocabulary from “Contributor Roles Defined” (CRediT) at https://credit.niso.org/contributor-roles-defined/. KJW was involved in conceptualization, formal analysis, investigation, methodology, software, validation, visualization, writing the original draft, and reviewing and editing. REB was involved in data curation, investigation, methodology, project administration, resources, and writing the original draft. BCS, MTG, JFL, TT and LT were involved in methodology, investigation, and resources. TB was involved in methodology, investigation, resources, and writing the original draft. KTG was involved in investigation, resources, and writing the original draft. MM and VMB were involved in resources and supervision. ARIL was involved with investigation, methodology, resources, supervision, and reviewing and editing. RWH and ILGN were involved with conceptualization, funding acquisition, investigation, project administration, resources, supervision, and reviewing and editing. JCDH was involved in conceptualization, data curation, funding acquisition, investigation, methodology, project administration, supervision, writing original draft, and reviewing and editing.

## References

1. Garalde DR, Snell EA, Jachimowicz D, Sipos B, Lloyd JH, Bruce M, Pantic N, Admassu T, James P, Warland A, Jordan M, Ciccone J, Serra S, Keenan J, Martin S, McNeill L, Wallace EJ, Jayasinghe L, Wright C, Blasco J, Young S, Brocklebank D, Juul S, Clarke J, Heron AJ, Turner DJ. 2018. Highly parallel direct RNA sequencing on an array of nanopores. Nat Methods 15:201–206.

2. Smith AM, Jain M, Mulroney L, Garalde DR, Akeson M. 2019. Reading canonical and modified nucleobases in 16S ribosomal RNA using nanopore native RNA sequencing. PLoS One 14:e0216709.

3. Workman RE, Tang AD, Tang PS, Jain M, Tyson JR, Razaghi R, Zuzarte PC, Gilpatrick T, Payne A, Quick J, Sadowski N, Holmes N, de Jesus JG, Jones KL, Soulette CM, Snutch TP, Loman N, Paten B, Loose M, Simpson JT, Olsen HE, Brooks AN, Akeson M, Timp W. 2019. Nanopore native RNA sequencing of a human poly(A) transcriptome. Nat Methods 16:1297–1305.

4. Zhang S, Li R, Zhang L, Chen S, Xie M, Yang L, Xia Y, Foyer CH, Zhao Z, Lam HM. 2020. New insights into Arabidopsis transcriptome complexity revealed by direct sequencing of native RNAs. Nucleic Acids Res 48:7700–7711.

5. Kim D, Lee JY, Yang JS, Kim JW, Kim VN, Chang H. 2020. The Architecture of SARS-CoV-2 Transcriptome. Cell 181:914–921 e10.

6. Viehweger A, Krautwurst S, Lamkiewicz K, Madhugiri R, Ziebuhr J, Holzer M, Marz M. 2019. Direct RNA nanopore sequencing of full-length coronavirus genomes provides novel insights into structural variants and enables modification analysis. Genome Res 29:1545–1554.

7. Bilinovich SM, Uhl KL, Lewis K, Soehnlen X, Williams M, Vogt D, Prokop JW, Campbell DB. 2020. Integrated RNA Sequencing Reveals Epigenetic Impacts of Diesel Particulate Matter Exposure in Human Cerebral Organoids. Dev Neurosci 42:195–207.

8. Ghedin E, Wang S, Spiro D, Caler E, Zhao Q, Crabtree J, Allen JE, Delcher AL, Guiliano DB, Miranda-Saavedra D, Angiuoli SV, Creasy T, Amedeo P, Haas B, El-Sayed NM, Wortman JR, Feldblyum T, Tallon L, Schatz M, Shumway M, Koo H, Salzberg SL, Schobel S, Pertea M, Pop M, White O, Barton GJ, Carlow CK, Crawford MJ, Daub J, Dimmic MW, Estes CF, Foster JM, Ganatra M, Gregory WF, Johnson NM, Jin J, Komuniecki R, Korf I, Kumar S, Laney S, Li BW, Li W, Lindblom TH, Lustigman S, Ma D, Maina CV, Martin DM, McCarter JP, McReynolds L, et al. 2007. Draft genome of the filarial nematode parasite Brugia malayi. Science 317:1756–60.

9. Tvedte ES, Gasser M, Sparklin BC, Michalski J, Hjelmen CE, Johnston JS, Zhao X, Bromley R, Tallon LJ, Sadzewicz L, Rasko DA, Dunning Hotopp JC. 2021. Comparison of long-read sequencing technologies in interrogating bacteria and fly genomes. G3 (Bethesda) 11.

10. Jones T, Federspiel NA, Chibana H, Dungan J, Kalman S, Magee BB, Newport G, Thorstenson YR, Agabian N, Magee PT, Davis RW, Scherer S. 2004. The diploid genome sequence of Candida albicans. Proc Natl Acad Sci U S A 101:7329–34.

11. Blattner FR, Plunkett G, 3rd, Bloch CA, Perna NT, Burland V, Riley M, Collado-Vides J, Glasner JD, Rode CK, Mayhew GF, Gregor J, Davis NW, Kirkpatrick HA, Goeden MA, Rose DJ, Mau B, Shao Y. 1997. The complete genome sequence of Escherichia coli K-12. Science 277:1453–62.

12. Bailey TL, Johnson J, Grant CE, Noble WS. 2015. The MEME Suite. Nucleic Acids Res 43:W39–49.

13. Harper JE, Miceli SM, Roberts RJ, Manley JL. 1990. Sequence specificity of the human mRNA N6-adenosine methylase in vitro. Nucleic Acids Res 18:5735–41.

14. Parker MT, Knop K, Sherwood AV, Schurch NJ, Mackinnon K, Gould PD, Hall AJ, Barton GJ, Simpson GG. 2020. Nanopore direct RNA sequencing maps the complexity of Arabidopsis mRNA processing and m(6)A modification. Elife 9.

15. Dominissini D, Moshitch-Moshkovitz S, Schwartz S, Salmon-Divon M, Ungar L, Osenberg S, Cesarkas K, Jacob-Hirsch J, Amariglio N, Kupiec M, Sorek R, Rechavi G. 2012. Topology of the human and mouse m6A RNA methylomes revealed by m6A-seq. Nature 485:201–6.

16. Schwartz S, Agarwala SD, Mumbach MR, Jovanovic M, Mertins P, Shishkin A, Tabach Y, Mikkelsen TS, Satija R, Ruvkun G, Carr SA, Lander ES, Fink GR, Regev A. 2013. High-resolution mapping reveals a conserved, widespread, dynamic mRNA methylation program in yeast meiosis. Cell 155:1409–21.

17. Bhattacharya T, Yan L, Crawford JM, Zaher H, Newton ILG, Hardy RW. 2022. Differential viral RNA methylation contributes to pathogen blocking in Wolbachia-colonized arthropods. PLoS Pathog 18:e1010393.

18. Bhattacharya T, Rice DW, Crawford JM, Hardy RW, Newton ILG. 2021. Evidence of Adaptive Evolution in Wolbachia-Regulated Gene DNMT2 and Its Role in the Dipteran Immune Response and Pathogen Blocking. Viruses 13.

19. Bhattacharya T, Newton ILG, Hardy RW. 2017. Wolbachia elevates host methyltransferase expression to block an RNA virus early during infection. PLoS Pathog 13:e1006427.

20. Bhattacharya T, Newton ILG, Hardy RW. 2020. Viral RNA is a target for Wolbachia-mediated pathogen blocking. PLoS Pathog 16:e1008513.

21. Chung M, Teigen L, Liu H, Libro S, Shetty A, Kumar N, Zhao X, Bromley RE, Tallon LJ, Sadzewicz L, Fraser CM, Rasko DA, Filler SG, Foster JM, Michalski ML, Bruno VM, Dunning Hotopp JC. 2018. Targeted enrichment outperforms other enrichment techniques and enables more multi-species RNA-Seq analyses. Sci Rep 8:13377.

22. Tvedte ES, Gasser M, Zhao X, Tallon LJ, Sadzewicz L, Bromley RE, Chung M, Mattick J, Sparklin BC, Dunning Hotopp JC. 2022. Accumulation of endosymbiont genomes in an insect autosome followed by endosymbiont replacement. Curr Biol 32:2786–2795 e5.

23. Li H. 2018. Minimap2: pairwise alignment for nucleotide sequences. Bioinformatics 34:3094–3100.

24. Danecek P, Bonfield JK, Liddle J, Marshall J, Ohan V, Pollard MO, Whitwham A, Keane T, McCarthy SA, Davies RM, Li H. 2021. Twelve years of SAMtools and BCFtools. Gigascience 10.

25. Shen W, Le S, Li Y, Hu F. 2016. SeqKit: A Cross-Platform and Ultrafast Toolkit for FASTA/Q File Manipulation. PLoS One 11:e0163962.

